# Evaluating the Biocontrol Potential of Pythium oligandrum and Serratia proteamaculans in Controlling Phytophthora plurivora in European beech (Fagus sylvatica)

**DOI:** 10.1101/2024.10.17.618890

**Authors:** Bekele Gelena Kelbessa, Anton Samuelsson, Farideh Ghadamgahi, Johanna Witzell, Stephen C. Whisson, Rodomiro Ortiz, Laura J. Grenville-Briggs, Ramesh R. Vetukuri

## Abstract

In recent years, the plant pathogen *Phytophthora plurivora* has caused severe damage to beech forests in many European countries, including Sweden. In many affected areas, few protective measures are in place due to fears about potential negative impacts on the ecosystem or public health. The research presented in this article assesses the biocontrol potential of the oomycete *Pythium oligandrum* and the bacterium *Serratia proteamaculans* against *P. plurivora*, with experiments performed under both *in vitro* and greenhouse conditions, which could represent a step forward in developing a safe treatment for European beech forests. The *in vitro* results revealed that *P. oligandrum* and *S. proteamaculans* significantly inhibited pathogen growth and stimulated a shift in the hyphal growth pattern towards shorter, branched hyphae with many hyphal swellings and thickened cell walls. The experiments conducted in greenhouses showed that treating three-month-old beech seedlings with *P. oligandrum* and *S. proteamaculans* counteracts the *P. plurivora* pathogen and reduces disease symptoms on the aerial and underground parts of the plant. GC-MS analysis detected the volatile organic compounds alpha-pinene, 2,5-dimethyl-pyrazine, and 3-methyl-1-butanol from *S. proteamaculans;* these compounds have the potential to inhibit pathogen growth. The disease suppression demonstrated by these biocontrol agents could thus be related to the synergistic effect of competition for nutrients with the secretion of hydrolytic enzymes and VOCs; moreover, induced systemic resistance may be triggered under greenhouse conditions. In conclusion, using *P. oligandrum* and *S. proteamaculans* could represent an environmentally friendly strategy for effectively controlling the disease caused by *P. plurivora* in beech.

## Introduction

European beech (*Fagus sylvatica*) is the most competitive tree species in both Western and Central Europe, and the mountainous regions of Eastern and Southern Europe, since it is highly shade tolerant and able to adapt to different environmental conditions ^1–6^. The presence of this tree species in natural and production forests is important both for the wood-processing industry and on a landscape ecology level ^7,8^. However, the gradual decline of beech populations across many European countries^9,10^. Among various biotic factors, responsible for this decline, the presence of *Phytophthora* spp. has emerged as an important causal agent in the last decade ^7^.

Although more than 17 *Phytophthora* species have been associated with beech forests ^11,12^, the three predominant species, which cause extensive damage to beech in many European countries are *Phytophthora plurivora*, *P. cactorum* and *P. cambivora* ^13–15^. These pathogens mainly attack fine roots, collars, and bark to cause tyloses in large xylem vessels, eventually blocking water and nutrient supply, and – in some cases – resulting in the death of the tree ^14,16,17^. Climate change is modifying weather patterns; for instance, excessive rainfall is now more commonly followed by extreme drought, a progression that exacerbates diseases caused by *Phytophthora* species, by promoting soil-borne zoospore production and stress in plants ^14^. In addition, global transport and the international plant trade mean that there are clear opportunities for the rapid spread of invasive species, such as *Phytophthora* spp., outside of their original geographical habitats ^18–20^, which can strongly affect the survival of native plant species and destabilize forest ecosystems ^21,22^.

In southern Sweden, the dominant *Phytophthora* species (*P. plurivora*, *P. cactorum* and *P. cambivora*) reported from other European countries also threaten several tree species, including *Fagus* spp., *Quercus* spp., *Alnus* spp., and *Pinus sylvestris*, all of which are important for the Swedish bio-economy ^22^. *P. plurivora* is a hemibiotrophic pathogen that seriously affects the health of young and mature beech trees by causing collar rot, bark canker, and crown dieback, and has become a major problem in southern Sweden ^22,23^. Effective control measures are therefore needed to combat the irreversible loss of beech trees to this aggressive pathogen. Unfortunately, the only viable strategy for combating this pathogen is the removal of infected trees, as commonly used synthetic fungicides may not be effective against *Phytophthora* spp. because they neither synthesize chitin nor ergosterol ^24^. In addition, chemical fungicides do not represent a viable option for disease control in natural ecosystems because this approach can harm human health and the environment. In recent years, low-risk plant protection products (PPPs) such as phosphite have been sought as a substitute for the more environmentally harmful synthetic pesticides to combat oomycetes and fungal diseases ^25^. However, the approval process for phosphite, which is recognized in many European countries for its potential to protect plants against diseases, including those caused by forestry pathogens, has not yet been completed in Sweden.

Furthermore, the use of this compound reduces the growth of *Phytophthora* pathogens without killing them, a characteristic, which is associated with the risk of resistance development if the compound is used excessively ^26^. In the last two decades, the use of plant-associated microbes, such as rhizobacteria, true fungi (*Trichoderma* spp.), and some *Pythium* spp., as biocontrol agents has emerged as a more sustainable alternative when compared to the application of synthetic fungicides to suppress plant diseases ^27–29^. Of the various *Pythium* species, *Pythium oligandrum*, which colonizes the root ecosystem of many plant species (e.g., tomato, pepper, and cucumber), is known to be a parasite of different fungal and oomycete pathogens such as *Botrytis cinerea*, *Fusarium oxysporum*, and *Phytophthora infestans* ^24,30,31^. *Py. oligandrum* utilizes direct (e.g., mycoparasitism and the production of antimicrobial compounds) and indirect (e.g., modulation of plant defense responses, possibly leading to the induction of systemic resistance) mechanisms to combat plant pathogens ^31,32^ and is unlikely to cause adverse negative effects on the surrounding plant and soil microbiomes. ^33^For example, *Py. oligandrum* can parasitize fungus, inhibiting pathogen growth by secreting enzymes involved in the degradation of the cell wall of fungal or oomycete prey species. It has also been reported that some species of the bacterial genus *Serratia* are associated with plants and can – in addition to improving plant growth – protect plants from pathogenic microbes ^34–36^. For example, it has been reported that *S. proteamaculans*, which can be isolated from the rhizosphere of naturally growing *Equisetum* plants, controls plant diseases caused by *Verticillium dahliae, Rhizoctonia solani*, and *Phytophthora colocasiae* ^37,38^. Although *P. oligandrum* and *S. proteamaculans* have been reported to suppress various soil-borne fungal and oomycete pathogens of herbaceous plants, the inhibitory effects of these organisms against *P. plurivora*, which infects woody beech trees, have not yet been investigated. Hence, this work aimed to evaluate the biocontrol potential of *P. oligandrum* and *S. proteamaculans* in reducing the disease associated with *P. plurivora* among beech plants under both *in vitro* and greenhouse conditions.

## Materials and Methods

### Microbial strains and culture conditions

*P. plurivora* AV1007 ^39^ was used in this study. The biocontrol agents tested in this study, a bacterial strain (*S. proteamaculans* strain S4 ^37^) and an oomycete strain (*Py*. *oligandrum* CBS530.74 ^40^) were provided by the Department of Plant Protection Biology, Swedish University of Agricultural Sciences (Alnarp, Sweden). Both *P. plurivora* and *Py. oligandrum* were maintained on V8 agar (CaCO_3_ 1 gL^-1^; agar 15 g L^-1^; V8 vegetable juice, 200 mL L^-1^) at 18 °C ^39^, while the culture of *S. proteamaculans* was maintained on Tryptic Soy Agar (TSA) (Sigma-Aldrich, St. Louis, MO, USA) at 8 °C.

### P. oligandrum inoculum

To obtain *P. oligandrum* oospores as inoculum, three mycelial plugs were incubated in 1l liquid V8 medium on a rotary shaker for five days at 18 °C as described in ^41^. The resulting cultures were homogenized in a blender for 1 minute, after which the oospores were separated from the mycelial debris by centrifugation at 5,000 × g for 10 minutes. The resulting pellet was resuspended in sterilized distilled H_2_O, mixed, and centrifuged again. These washing steps were performed three times. Following the final washing step, the oospores were resuspended in sterilized distilled H_2_O, after which the concentration was measured using a Fuchs-Rosenthal hemocytometer (Karl Hecht, Germany) and adjusted to a final concentration of 30 × 10^3^ oospores/mL for use in the experiments.

### S. proteamaculans inoculum

*S. proteamaculans* was cultured overnight in half-strength tryptone-soya broth (TSB) on a rotary shaker at 28 °C. The concentration was determined based on serial dilutions of the suspensions in TSA at several time points, with the absorbance of the suspension measured at 600 nm using a spectrophotometer (Eppendorf AG, Hamburg, Germany). Subsequently, the concentration of colony-forming units (CFUs) at each time point was determined, and the final concentration was adjusted to 10^6^ CFU/mL and used in this study.

### P. plurivora zoospores

For the collection of *P. plurivora* zoospores, water from hydroponically grown beech seedlings (2 seedlings L^-1^) was filtered through a 40 µm filter, with 20 mL of the filtered beech water added to an empty 9 cm diameter Petri dish. Plugs from the edges of the 14-day-old actively growing *P. plurivora* cultures on V8 agar were cut and then aseptically transferred to a Petri dish containing filtered beech water (6 plugs per empty Petri dish). Cultures were maintained in growth chambers at 23°C with 85 % RH and a photoperiod of 16 hours light and 8 hours darkness, and the water was replaced with sterilized distilled H_2_O after 48 hours. The number of sporangia was quantified under a microscope after 72 hours. The plates were then incubated at 4 °C in the refrigerator for 1 hour, after which the plates were placed back in the growth chamber for 2 hours to stimulate the release of zoospores. The H_2_O containing the zoospores was then carefully transferred from the Petri dishes to 50-mL falcon tubes and centrifuged at low speed (2500 × g) to avoid encysting the zoospores. The supernatant was carefully discarded, and the liquid remaining at the bottom of the falcon tubes, which contained the zoospores, was collected in a common tube and gently stirred to ensure homogeneity before extracting the zoospores. The zoospore concentration was then measured with a hemocytometer, with the final concentration adjusted to 5 × 10^5^ zoospores/mL for use in the study.

### Dual culture assay

The inhibitory effect of *Py. oligandrum* on *P. plurivora* growth was investigated via a time-course dual culture assay using cornmeal agar (CMA) (Sigma Aldrich) plates. Briefly, 5 mm mycelial plugs, taken from the edges of actively growing colonies of both oomycetes, were inoculated at opposite sides of a 9 cm diameter Petri dish and allowed to grow in the dark at 18°C for 20 days. In addition, both oomycetes were inoculated together in a thin layer of CMA covering the bottom of Petri dishes, and mycelial samples from the interaction zone were collected 5, 6, and 7 days later for microscopic visualization. In these dual culture assays, two treatments were performed with three replicates per treatment: 1 – *P. plurivora* alone; and 2-*Py. oligandrum* + *P. plurivora*.

### Py. oligandrum filtrate assay

We also assessed how the addition of *Py. oligandrum* filtrate affects *P. plurivora* growth. In this experiment, a 5 mm mycelial plug was taken from an actively growing colony of *Py. oligandrum* and transferred to a 50 mL Falcon tube containing 5 mL of V8 broth; this tube was then kept in the dark at 18°C for 72 hours. After the incubation period, the liquid from the Falcon tube was filtered through a 0.4 µm filter. Subsequently, 1 mL of the filtrate from the culture was carefully spread onto CMA plates and allowed to dry for 30 min before a 5 mm plug of *P. plurivora* was inoculated into the center of the 9 cm diameter Petri dish. This assay also included two treatments with three replicates: 1 – *P. plurivora* alone; and 2 – *Py. oligandrum* filtrate + *P. plurivora*. The control treatment was treated with 1 mL of filtered liquid V8. The plates were incubated at 18°C for 14 days. At the end of the incubation time, the inhibition of mycelial growth was assessed as follows [Radial inhibition (%) = ((R1-R2)/R1 ×100), where R1 is the radial growth of the pathogen in the absence of *Py. oligandrum* filtrate and R2 is the radial growth of the pathogen in the presence of *Py. oligandrum* filtrate ^35^.

### S. proteamaculans antagonism assay

The antagonistic effect of *S. proteamaculans* against *P. plurivora* was investigated on a CMA plate. In this assay, 5 mm plugs were collected from two-week-old *P. plurivora* cultures growing on V8 agar, and then inoculated into the center of a 9 cm diameter Petri dish; the overnight cultured suspension of *S. proteamaculans* was then streaked on opposing sides of the plates in 4 cm lines at an equal distance from the plug. This assay included two treatments with three replicates: 1 – *P. plurivora* alone; and 2 – *S. proteamaculans* + *P. plurivora*. The plates were placed in the dark at 24°C for 28 days. At the end of the incubation period, the inhibition of radial growth was measured as follows [Radial inhibition (%) = ((R1-R2)/R1 ×100), where R1 is the radial growth of the pathogen in the absence of *S. proteamaculans* and R2 is the radial growth of the pathogen in the presence of *S. proteamaculans* ^35^. The interaction between *S. proteamaculans* and *P. plurivora* was also studied on a thin layer of CMA, and mycelial samples from the interaction zone were collected 6, 10, and 20 days later for microscopic visualization. These dual culture experiments also included two treatments with three replicates: 1 – *P. plurivora* alone; and 2 – *S. proteamaculans* + *P. plurivora*.

### S. proteamaculans filtrate assay

To investigate the antagonistic activity of a cell-free filtrate from *S. proteamaculans* against *P. plurivora*, the bacterium was inoculated into 20 mL of LB in 50 mL Falcon tubes and incubated for 16 hours at 18°C with constant shaking at 220 rpm. After incubation, the supernatants were collected by centrifugation at 4200 × g for 10 min at room temperature and filtered through a 0.22 μm membrane (Millipore, Billerica, MA). The culture filtrates were then added to pre-cooled CMA medium at a final concentration of 10 % (v/v), and 20 mL of the prepared CMA medium was added to 9 cm diameter Petri dishes. The control plates contained only CMA. Next, 5 mm plugs collected from a 14-day-old culture of *P. plurivora* were inoculated into the center of the Petri dish and incubated at 18°C for 14 days. This assay contained two treatments with three replicates per treatment: 1 – *P. plurivora* alone; and 2 – S4 filtrate + *P. plurivora*. At the end of the incubation period, percentage pathogen growth inhibition was calculated as follows [Radial inhibition (%) = ((R1-R2)/R1×100) ^35^.

### GC-MS detection of volatile organic compounds (VOCs)

As we were interested in *S. proteamaculans*, the volatile organic compounds (VOCs) produced by this bacterium were detected by gas chromatography coupled to mass spectrometry (GC-MS). A single colony of *S. proteamaculans* was cultured in 50 mL nutrient broth for 15-16 hours in a shaking incubator at 28°C and 200 rpm according to the method described by Revadi et al. ^42^. The bacterial suspension was transferred to a 500 mL wash bottle under sterile conditions. The bacterial headspace was collected by pushing 0.1 L/min of charcoal-filtered clean air through the suspension onto an adsorption filter (Super Q trap 80/100 mesh; Alltech, Deerfield, IL, USA) attached to the outlet of the wash bottle for approximately three hours. The adsorption filter was then rinsed with 0.5 mL hexane and the sample was used for GC-MS analysis (6890 GC and 5975 MS; Agilent Technologies, Santa Clara, CA, USA). Next, 2 μL of the sample was injected into an HP-5MS column (30 m × 0.25 mm × 0.25 μm film thickness; J&W Scientific, Folsom, CA, USA) programmed for 37 min with the following temperature settings: initial temperature of 50°C (2 min) followed by an 8°C/min increase to a final temperature of 275°C (held for 10 min). Peaks were identified by injecting standard reference compounds and comparing the results with Kovát’s retention indices and mass spectrum from the NIST reference library (Agilent) ^43^.

### Greenhouse trials

A greenhouse trial was carried out to evaluate the biocontrol effects of *P. oligandrum* and *S. proteamaculans* in counteracting the adverse impacts of *P. plurivora* infection among beech seedlings. Briefly, similar-sized, three-month-old seedlings of the beech variety ‘Gottåsa’ were sourced from a local nursery (Ranviks Trädgård, Båstad, Sweden), after which the roots were washed with deionized water and checked for macroscopic damage. Healthy beech seedlings were then planted in a 1 L pot filled with sterilized compost (Krukvåxtjord Lera and Kisel, Sweden) and kept under greenhouse conditions of 25/15°C (day/night) with 70% relative humidity and a 16:8 light:dark photoperiod. The beech seedlings were treated with *S. proteamaculans* at a final concentration of 10^6^ colony-forming units (CFUs) per mL. Treatment with *P. oligandrum* was carried out using an oospore suspension with a concentration of 30×10^3^ oospores/mL. For each treatment, 50 mL of the *P. oligandrum* suspension or *S. proteamaculans* was applied topically to the soil around the base of the plant. A medium devoid of any added microbes was used for the untreated control. After 24 hours, the plants were infected with *P. plurivora* zoospores by pouring 50 mL/pot (5 × 10^5^ zoospores/mL) into the soil around the base of the plant. Subsequently, all of the plants were placed into growth chambers for 30 days at 21/18°C (day/night), 65% relative humidity, and a 16:8 light:dark photoperiod. The pots under various treatments were rotated and watered twice a week and the aerial parts of the plants were monitored during the experiment.

At the end of the four-week experimental period, all of the plants were removed from the growth chamber so that the yellowing/wilting of leaves and macroscopic root damage could be assessed by visual observation for each individual plant. The estimated yellowing/wilting of leaves and root damage of the beech plants were scored using one of the following rating categories (0-5 scale): 0 = no visible yellowing of leaves/root damage visible; 1 = 1-20% yellowing of leaves/root damage; 2 = 21-40% yellowing of leaves/root damage; 3 = 41-60% yellowing of leaves/root damage; 4 = 61-80% yellowing of leaves/root damage, and 5 = 81-100% yellowing of leaves/root damage. Subsequently, the disease severity index (DSI) for the yellowing or wilting of leaves and damage to the main and lateral roots was calculated for each disease rating as follows [(DSI = (sum of numerical ratings) / (number of plants examined × maximum disease scale) × 100)] ^44^. In addition, the length of the dark lesion-cankers associated with *P. plurivora* and observed on the taproot was measured (mm). After disease assessment, each plant was cut into two parts, i.e., shoots and roots, which were placed into separate wax paper bags and dried at 70°C for 24 hours; after sufficient drying, the dry weight of the shoots and roots was measured (g). A total of six treatments were included in the greenhouse trial: 1-Water control; 2-*P. plurivora* alone; 3-*S. proteamaculans* alone; 4-*P. oligandrum* alone; 5-*S. proteamaculans* + *P. plurivora*; 6-*P. oligandrum* + *P. plurivora*. Treatments were conducted in a completely randomized design which included six replicates.

### Statistical analysis

The data obtained from the *in vitro* antagonistic assays and greenhouse experiments were checked for normality using Shapiro-Wilk’s and Levenés tests (*P > 0.05*). Afterward, the statistical significance of differences between the mean values associated with various treatments was analyzed using Duncan’s multiple range test in R software (version 4.2.0); the threshold for statistical significance was set as *P≤0.05*.

## Results

### Antagonistic assays

The *in vitro* dual culture of *Py. oligandrum* and *P. plurivora* on CMA plates demonstrated some degree of tolerance in occupying the same growth space, with no clear inhibition observed during the first four days of co-inoculation. However, the presence of *Py. oligandrum* significantly suppressed the dense vegetative growth of *P. plurivora* over 20 days (63.56%) when compared to the untreated control (Fig. 1A and B). In comparison, the mycelial growth of *P. plurivora* was not significantly suppressed when only the filtrate of this antagonistic oomycete was added to the CMA plates; suggesting that competition for nutrients, rather than the bioactive compounds present in the filtrate, represents the main mechanism through which *Py. oligandrum* inhibits *P. plurivora* growth (Fig. 1C). Appressorium-like structures and coils of *Py. oligandrum* were observed during hyphal interactions, and this appeared to suppress the mycelial growth of *P. plurivora* (Fig. 2A and B). However, most *Py. oligandrum* hyphae were devoid of cytoplasm after 20 days, which is likely the result of nutrient starvation (Fig. 2C).

**Figure 1:**
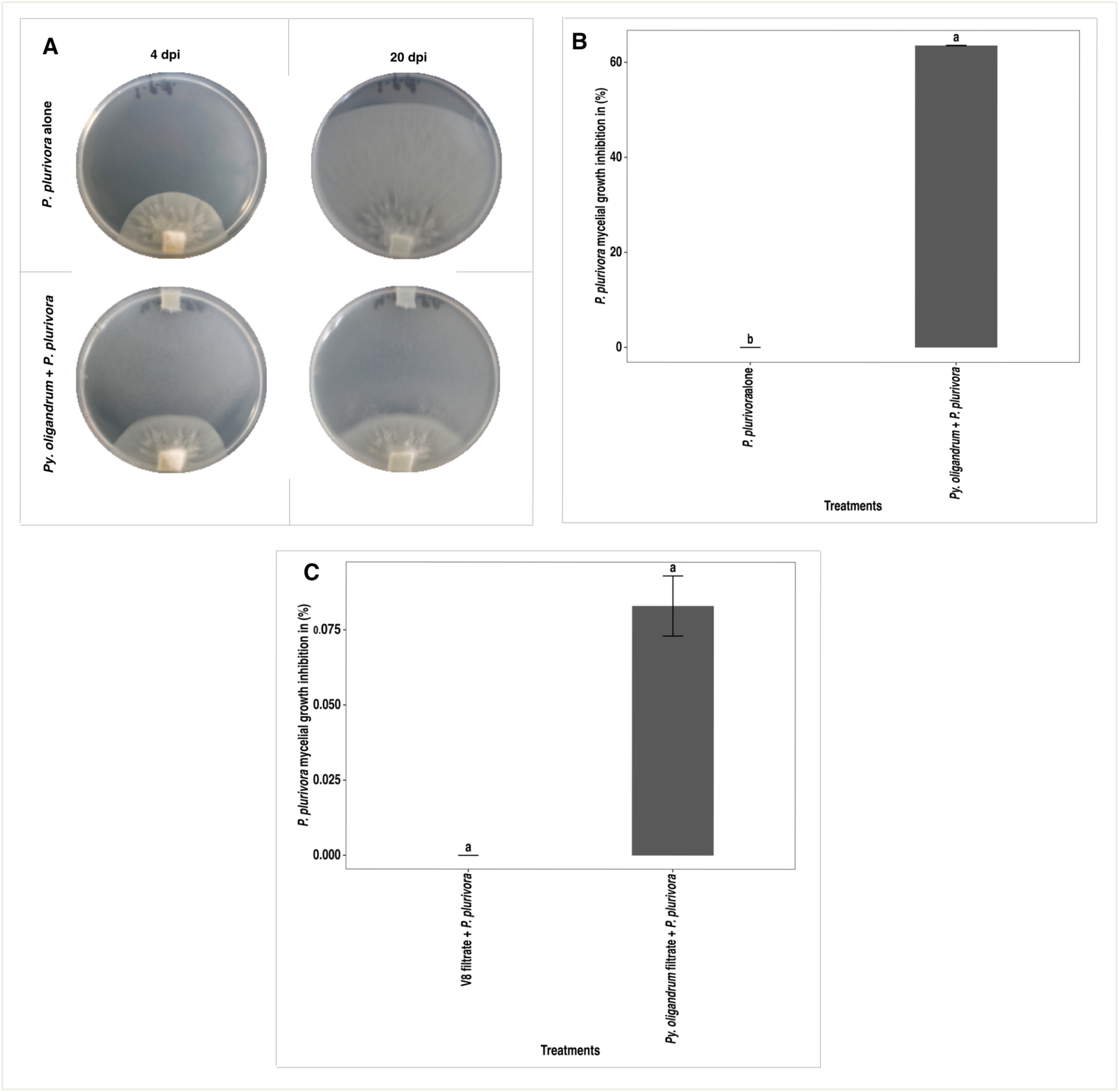
Antagonistic activity of *Py. oligandrum* on mycelial growth of *P. plurivora* observed during *in vitro* experiments. (A) Representative image of the untreated control (top row) and the interaction between *P. plurivora* and *Py. oligandrum* (bottom row) after four and 20 days of growth at 18°C in the dark on CMA plates. (B) Percentage inhibition of *P. plurivora* mycelial growth after 20 days of co-culturing with *Py. oligandrum*. (C) Effect of *Py. oligandrum* filtrate on pathogen growth inhibition (%). Data are presented as the mean value ± standard deviation, while mean values with different letters indicate a significant difference between treatments at *P≤ 0.05* according to Duncan’s multiple range test.

**Figure 2:**
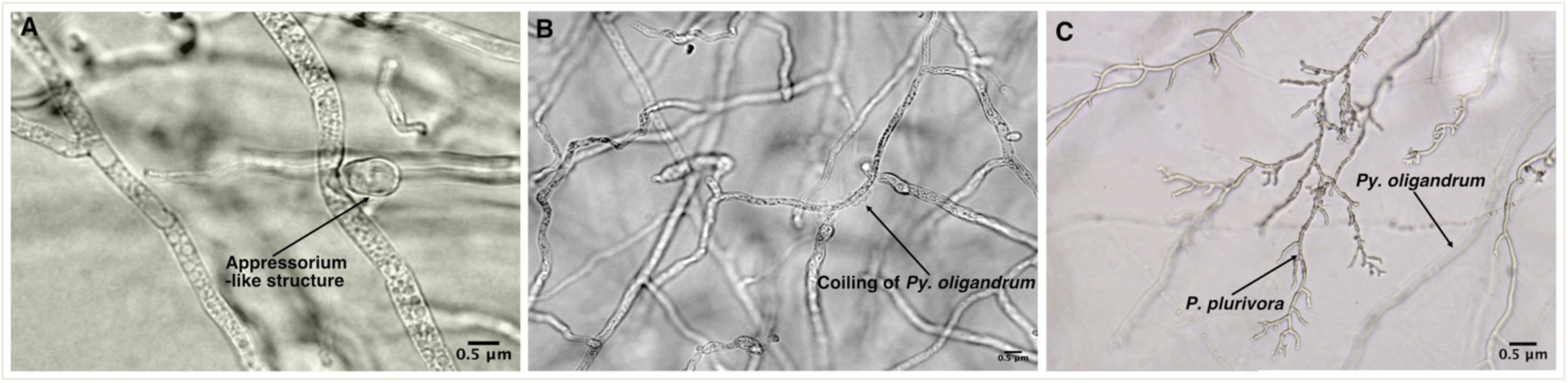
Hyphal interactions between *Py. oligandrum* and *P. plurivora* on thin-layered CMA plates, with the presented images captured using a Leica DMLB fluorescence microscope. (A) An appressorium-like structure of *Py. oligandrum* pressing against the hyphae of *P. plurivora* after four days of co-inoculation (400× magnification). (B) The coiling of *Py. oligandrum* hyphae around the hyphae of *P. plurivora*, resulting in erratic growth and hyphal swelling of the latter; the image was captured after six days of co-inoculation at 200× magnification. (C) Hyphal interactions between *Py. oligandrum* (bottom) and *P. plurivora* (top), with the former showing an empty cytoplasm and the latter a coralloid branching pattern after 20 days of co-inoculation (100 x magnification).

### *S. proteamaculans* antagonistic assay

Co-inoculation with *S. proteamaculans* significantly inhibited the mycelial growth and altered the hyphal morphology of *P. plurivora* (47.5%) (Fig. 3A and B). Hyphal growth in the presence of *S. proteamaculans* was characterized by secondary growth patterns with many aerial structures and hyphal swellings (e.g., globulose, pyriform, coralloid, and arbuscular structures); in contrast, the untreated control (Fig.4B, C, and D) maintained a long, filamentous hyphal growth pattern. However, older hyphae closer to the initial plugs in the control plates did show some branched and irregular patterns, which could possibly be explained by local adaptation of the hyphae to low nutrient content. Microscopic visualization of the interaction plates also revealed the presence of hyphal clusters and gelatinous droplets (Fig. 4A). Similarly, *P. plurivora* growth was significantly inhibited following the addition of the cell-free filtrate of *S. proteamaculans* when compared to the untreated control (Fig. 3C). The profile of VOCs in the headspace of *S. proteamaculans* cultures was analyzed by GC-MS analysis, with the results identifying 11 VOCs representing distinct chemical classes, including one alkane, five organic compounds, three alcohols, one aromatic compound, and one methyl ketone (Table 1).

**Figure 3.**
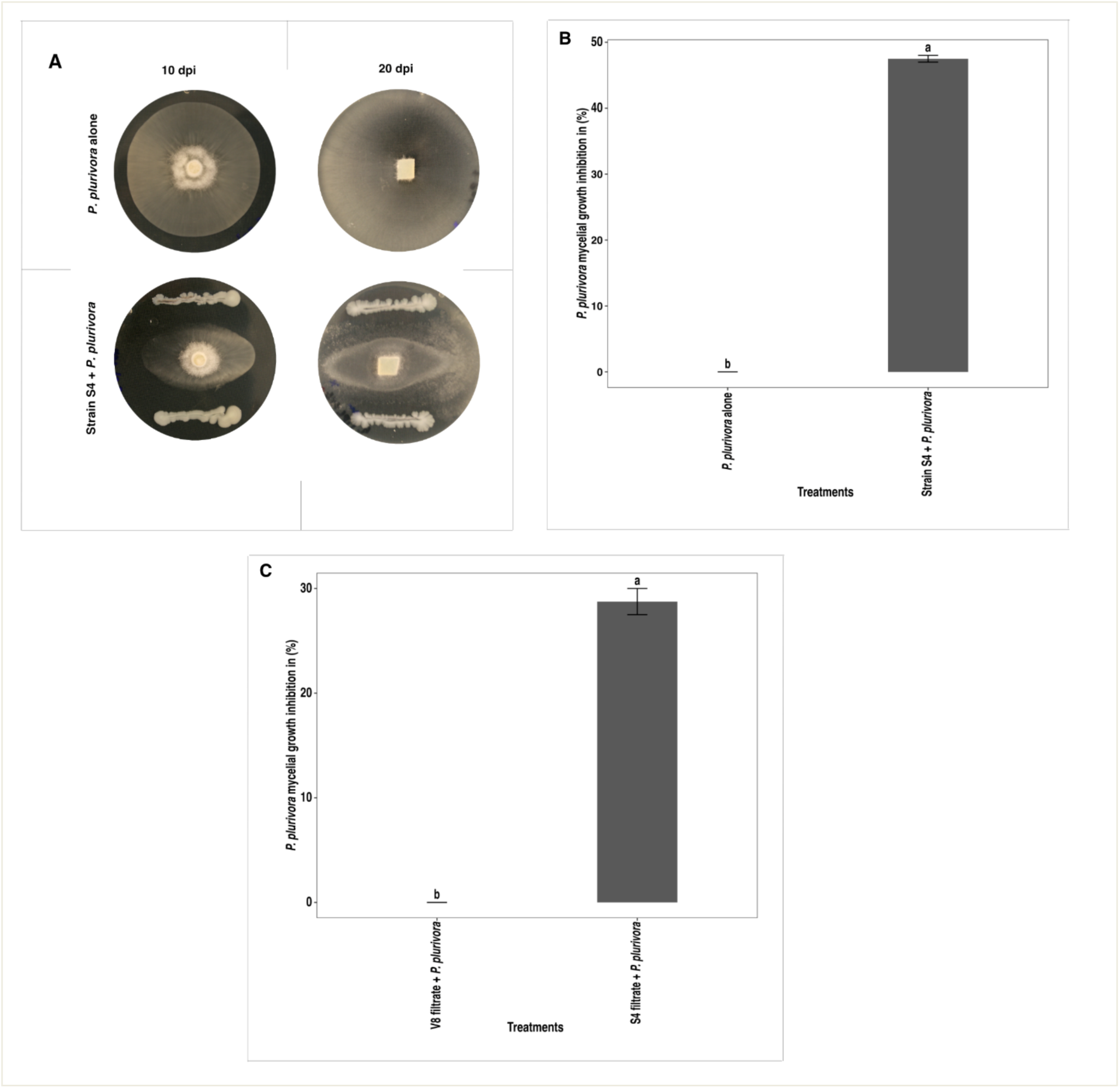
The antagonistic activity of *S. proteamaculans* against *P. plurivora* observed during *in vitro* experiments. (A) Image of the untreated control (top row) and the interaction between *S. proteamaculans* and *P. plurivora* (bottom row) after 10 and 20 days of growth at 24°C in the dark on CMA plates. (B) Inhibition (%) of *P. plurivora* mycelial growth after 14 days of co-culturing with *S. proteamaculans*. (C) Effect of the cell-free filtrate of *S. proteamaculans* on the inhibition of pathogen growth (%). Data are presented as mean values ± standard deviation, while mean values with different letters indicate significant between-treatment differences at *P≤ 0.05* according to Duncan’s multiple range test.

**Figure 4:**
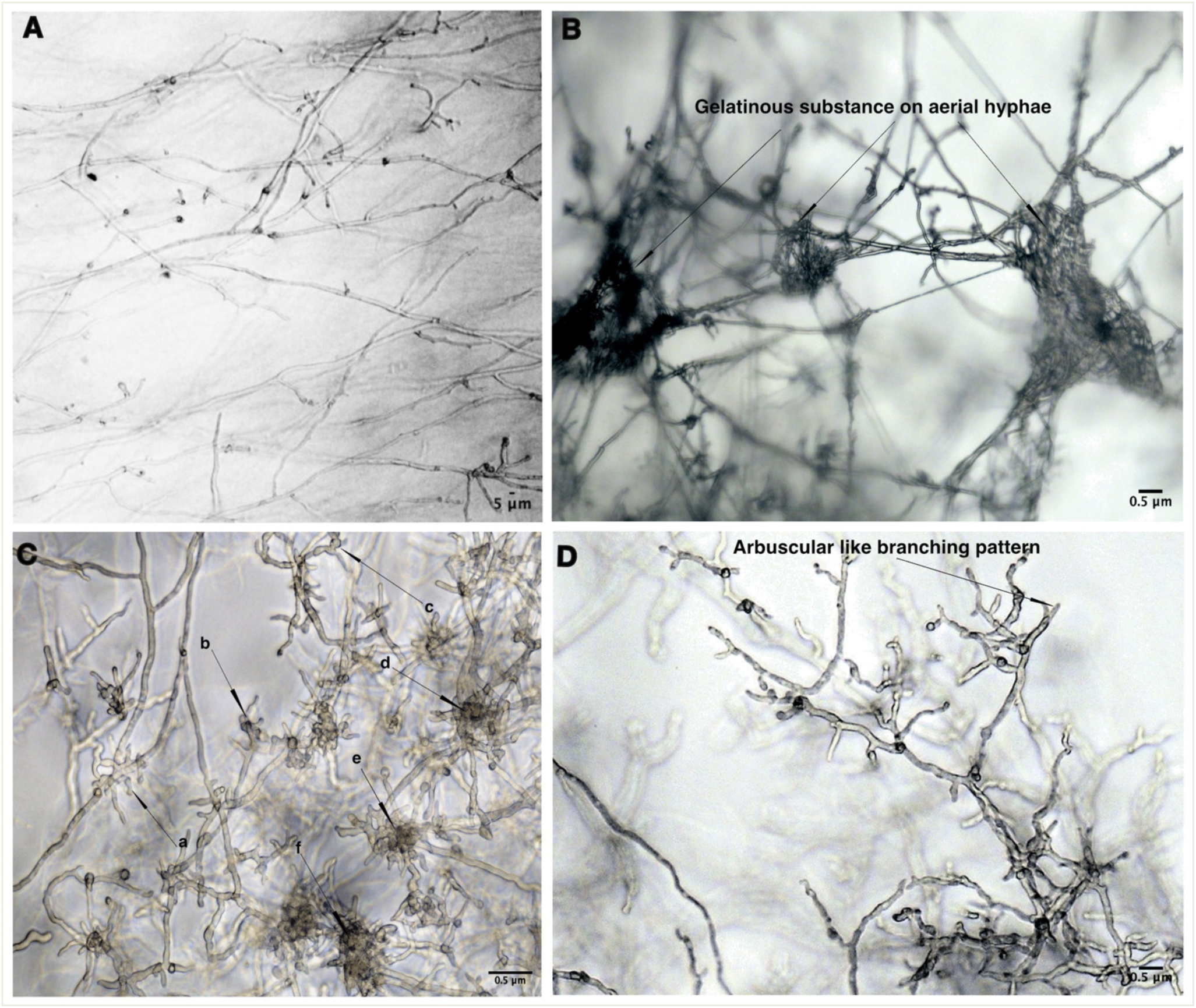
Hyphal interactions between *S. proteamaculans* and *P. plurivora* on thin layer CMA plates, with pictures captured using a Leica DMLB fluorescence microscope. Images of (A) untreated control plates after 10 days of inoculation (100x magnification), (B) clumped aerial hyphae of *P. plurivora* with the gelatinous substance after six days of co-inoculation (100× magnification), (C) erratic branching patterns, hyphal swellings (a, b, c), and hyphal clusters (d, e, f) of *P. plurivora* after 10 days of co-inoculation (200× magnification), and (D) arbuscular-like branching patterns of *P. plurivora* hyphae after 20 days of co-inoculation (100x magnification).

**Table 1:** Volatile organic compounds detected in the headspace of *S. proteamaculans* cultures.

| Chemical class | Compound | Chemical formula | Retention time (min) | GC peak relative area |
| --- | --- | --- | --- | --- |
| Alkanes | Decane | C<sub>10</sub>H<sub>22</sub> | 9.12 | 0.00 ± 0.00 |
|  | alpha-Pinene | C<sub>10</sub>H<sub>16</sub> | 9.50 | 0.00 ± 0.00 |
|  | Dimethyl disulfide | C<sub>2</sub>H<sub>6</sub>S<sub>2</sub> | 10.56 | 0.12 ± 0.04 |
| Miscellaneous organic compounds | Dimethyl trisulfide | CH<sub>3</sub>SSSCH<sub>3</sub> | 16.55 | 0.06 ± 0.02 |
|  | 4-ethoxy-ethyl ester-benzoic acid | C<sub>11</sub>H<sub>14</sub>O<sub>3</sub> | 27.77 | 0.00 ± 0.00 |
|  | 2,5-dimethyl-Pyrazine | C<sub>6</sub>H<sub>8</sub>N<sub>2</sub> | 15.41 | 0.00 ± 0.00 |
| Alcohols | 1-butanol | C<sub>4</sub>H<sub>9</sub>OH | 11.74 | 0.01 ± 0.00 |
|  | 3-methyl-1-butanol | C<sub>5</sub>H<sub>12</sub>O | 13.10 | 0.17 ± 0.05 |
|  | 1-Decanol | C<sub>10</sub>H<sub>22</sub>O | 22.26 | 0.01 ± 0.00 |
| Aromatic compounds | p-Cymene (1-methyl-4-(1-methylethyl)-benzene) | OC<sub>10</sub>H<sub>14</sub> | 14.44 | 0.00 ± 0.00 |
| Methyl ketones | 3-hydroxy-2-butanone | (CH<sub>3</sub>CH(OH)COCH<sub>3</sub>)<sub>2</sub> | 14.58 | 0.01 ± 0.00 |

### Greenhouse trials

In the greenhouse trials, beech plants treated with either *Py. oligandrum* or *S. proteamaculans* showed few symptoms of *P. plurivora* root rot disease, i.e., taproot and lateral root rot and leaf discoloration, compared to plants treated with *P. plurivora* alone. The roots of most plants treated with *P. plurivora* alone were severely damaged and most leaves were discolored (wilted) (Fig. 5B and Fig. 6B, C, and D). In contrast, the lateral roots of a few plants treated with *S. proteamaculans* or *P. oligandrum* plus *P. plurivora* showed disease symptoms, while most other plants were visually healthy. This suggests that treatment with strain *S. proteamaculans* or *P. oligandrum* can suppress the disease caused by *P. plurivora* and reduce the effects on beech plants. However, the biocontrol potential of treatment with *S. proteamaculans* and *P. oligandrum* was only significant in suppressing the lateral root rot associated with *P. plurivora* disease, as leaf yellowing was similar in the beech plants treated with water, with *S. proteamaculans* alone, or with *P. oligandrum* alone. Hence, the results can only be considered as being indicative of the antagonistic effects of the tested biocontrol agents. Additionally, the dry mass (e.g., the dry weight of the roots) of the diseased plants was equivalent to that of the treated plants, although the dry weight of the shoots of treated plants was higher than that measured for the control plants (Fig. 6E and F). These results indicate that the beech plants did not grow much during the experiment, i.e., all of the biomass was present at the start of the experiment and very little additional biomass was produced.

**Figure 5:**
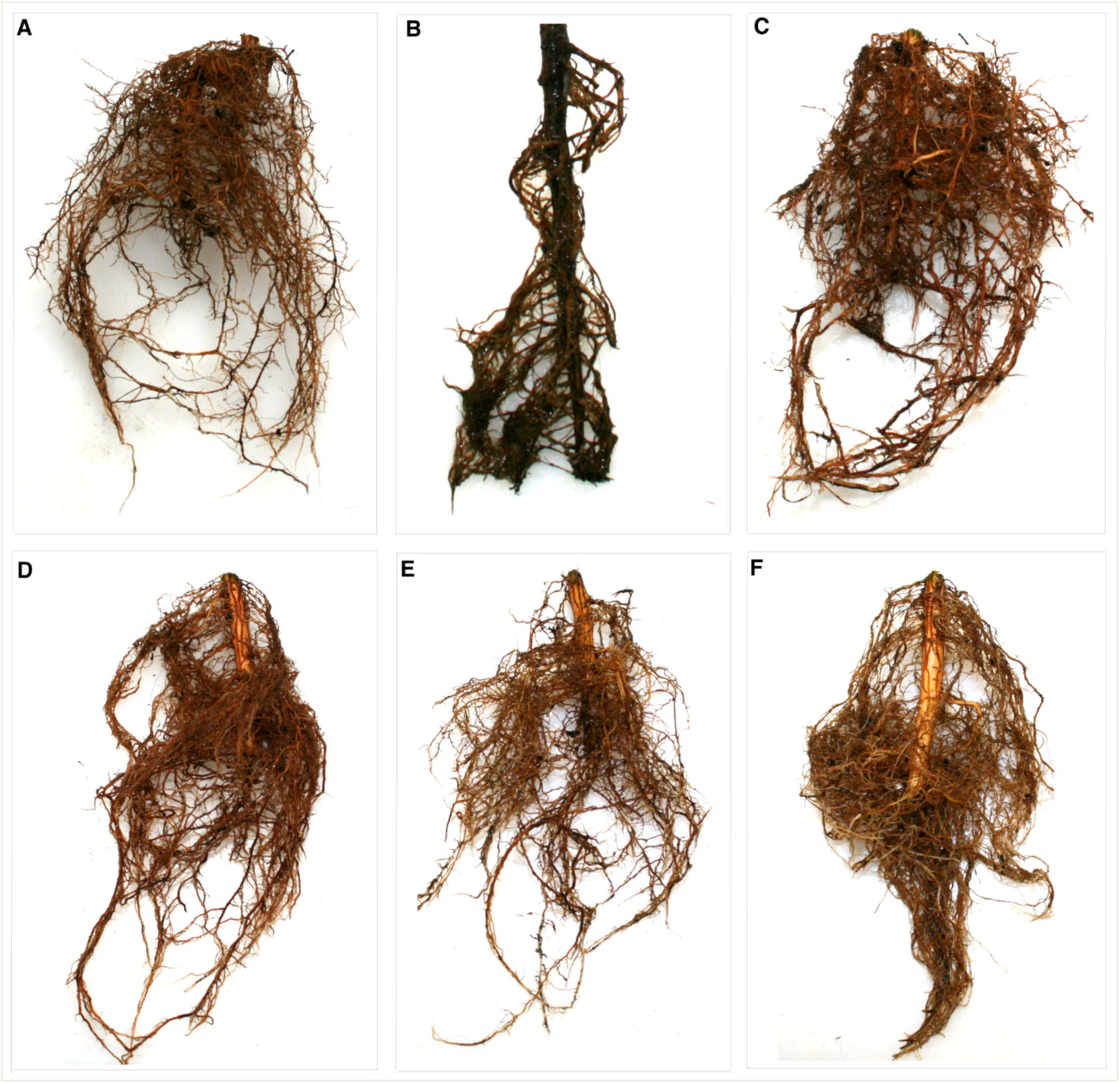
The results of the greenhouse biocontrol trial against *P. plurivora* in terms of root pictures. Results for (A) water control, (B) *P. plurivora* control, (C) *S. proteamaculans* control, (D) *P. oligandrum* control, (E) *S. proteamaculans* + *P. plurivora*, and (F) *P. oligandrum* + *P. plurivora*.

**Figure 6:**
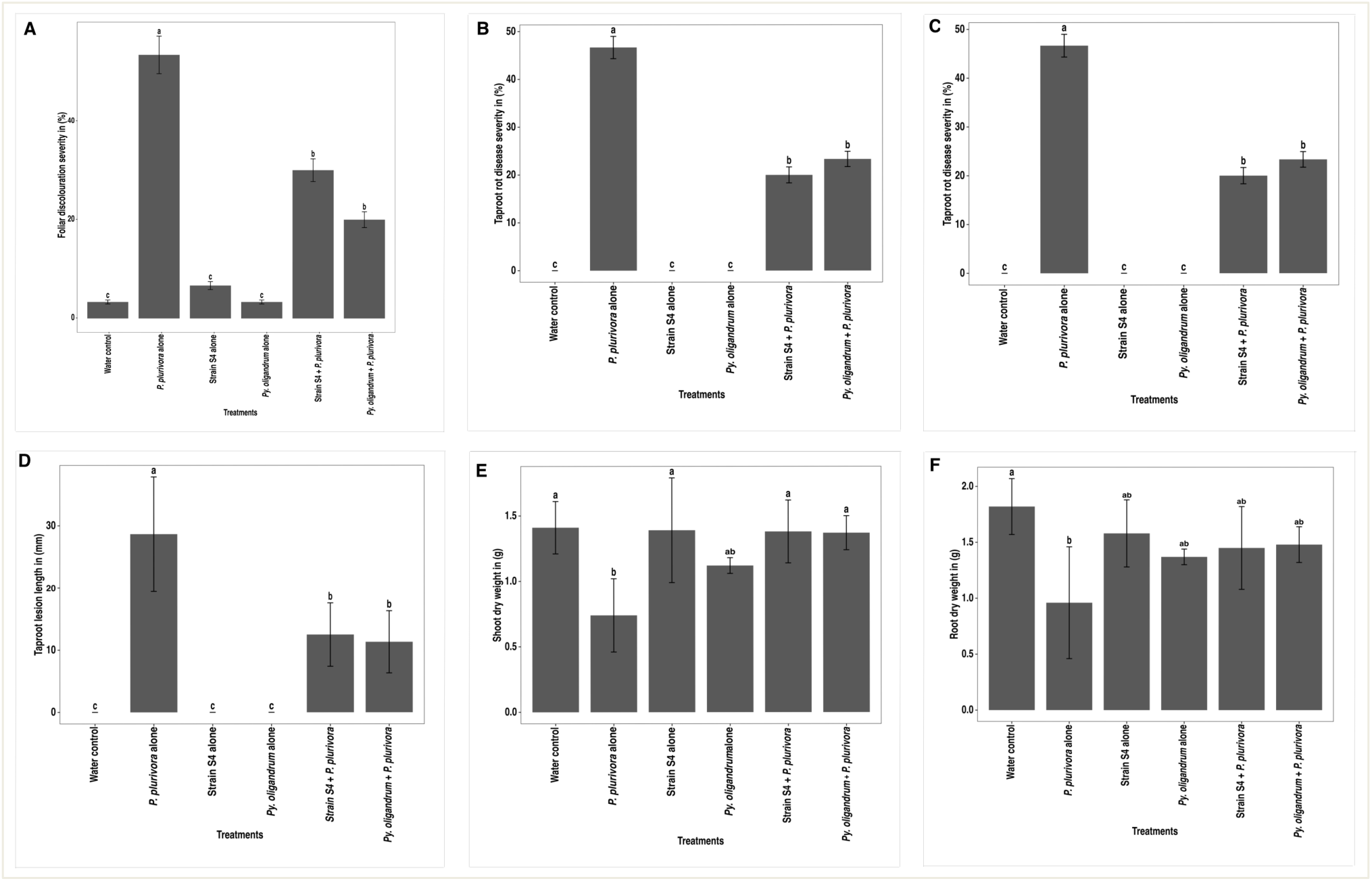
The efficacy of *S. proteamaculans* and *Py. oligandrum* in suppressing the above-and below-ground damage to beech plants associated with *P. plurivora* under greenhouse conditions. (A) Foliar disease severity (%) among beech plants. (B) Taproot disease severity (%) among beech plants. (C) Lateral root disease severity (%) among beech plants. (D) Length of taproot root infection (mm). (E) Shoot dry weight (g). (F) Root dry weight (g). Data are presented as mean values ± standard deviation, while mean values with different letters indicate statistically significant differences between groups at *P≤ 0.05* according to Duncan’s multiple range test.

## Discussion

The oomycete *Py. oligandrum* and the bacterium *S. proteamaculans* were evaluated under both *in vitro* and *in planta* conditions for biocontrol potential against *P. plurivora*. The results showed that *P. plurivora* growth was significantly inhibited *in vitro* in the presence of *Py. oligandrum*, and our data suggest that this effect may be due to direct mycoparasitic interaction between *Py. oligandrum* and *P. plurivora* hyphae as well as competition for nutrients and space. However, the culture filtrate from *Py. oligandrum* showed little effect on *P. plurivora* growth suggesting that any antibiotic substances secreted by this oomycete play a lesser role in directly suppressing pathogen growth, in this case, the growth of *P. plurivora*. The presence of *Py. oligandrum* also significantly reduced leaf wilting and root damage *in planta* when the results were compared to what was observed in untreated beech plants, i.e., clear signs of *P. plurivora* disease on the leaves and roots. In addition to mycoparasitic interactions and competition for nutrients and space, *Py. oligandrum* has also been reported to induce systemic resistance to legume pathogens without affecting mutual interactions ^45^.

*Py. oligandrum* has been reported to suppress oomycete and fungal pathogens via direct (e.g., secretion of antibiotic substances, mycoparasitism, and niche competition) and indirect mechanisms (e.g., triggering induced systemic resistance) ^45–48^. Parasitic coiling is associated with the production of cellulase, which enables *Py. oligandrum* to penetrate hyphae and effectively destroy pathogenic microbes; this has been cited as one of the most important mechanisms for suppressing various fungal and oomycete pathogens ^24,30,49,50^. Recently, Liang et al.^50^ showed that during mycoparasitic interaction with oomycetes or fungi, *P. oligandrum* regulates many carbohydrate-related gene families and degrades the components of the pathogen’s cell walls such as cellulose, and hemicellulose. In some cases, *Py. oligandrum* can produce an antimicrobial compound specific to the target pathogen, e.g., *Phytophthora megasperma;* in this case, the activity of *Py. oligandrum* can inhibit the growth of pathogen without degrading the cell wall^51^. *P. oligandrum* also probably stimulates the immune system of beech plants under greenhouse conditions by activating disease-resistance genes. Studies have shown that *Py. oligandrum* induces systemic resistance in the plant by activating many pathogen-associated R-family genes, stimulating subtilases associated with root symbionts, and regulating key plant hormones involved in stress responses ^24,33,52^.

*S. proteamaculans* also inhibited *P. plurivora* growth *in vitro*. This bacterium also suppressed the progression of the disease caused by *P. plurivora* on the above- and below-ground parts of beech plants under greenhouse conditions. Changes in the pathogen mycelium at the interaction zone could be related to degradation of the pathogen cell wall by specific enzymes secreted by the bacterial strain, or the production of organic compounds that are toxic to oomycetes by *S. proteamaculans*.

The results presented in our previous study ^38^ revealed that the different cell wall-degrading enzymes, including chitinase, cellulase, protease, and lipase, secreted by *S. proteamaculans* belong to the class of compounds that many plant growth-promoting rhizobacteria (PGPR) secrete to combat a broad spectrum of fungal and oomycete pathogens ^53,54^. However, the possibility that other antibiotic substances and VOCs are involved in the inhibitory activity of this strain cannot be ruled out. The bacterial filtrate assay confirmed that antibiotic substances released into the medium, rather than competition for nutrients, may play a major role in the *in vitro* inhibition of pathogen growth. The GC-MS analysis also detected various VOCs, including alpha-pinene, decane, dimethyl-disulfide, dimethyl-trisulfide, 2,5-dimethyl-pyrazine, 3-hydroxy-2-butanone, and 3-methyl-1-butanol, from cultures of this bacterial strain; these compounds, which could be potent inhibitors of *P. plurivora* mycelial growth, have also been found to be emitted by other *Serratia* strains ^55–57^.

Two VOCs, namely, 4-ethoxy-ethyl ester-benzoic acid and p-cymene (1-methyl-4-(1-methyl ethyl)-benzene), have not been detected from cultures of other *Serratia* strains, and may thus only be produced by *S. proteamaculans*. Previous reports have stated that the VOCs synthesized by rhizobacterial strains can protect plants against fungal, oomycete, and bacterial diseases; as such, these compounds could be used to control plant diseases ^58–60^. Numerous studies have highlighted how the VOCs synthesized by rhizobacterial strains (e.g., *Serratia*, *Pseudomonas*, and *Bacillus*) can inhibit pathogenic microbes and regulate plant growth activities. Thus, it is likely that the presence of *S. proteamaculans* can reduce the severity of the plant disease caused by *P. plurivora* via synergistic effects between different mechanisms such as the action of cell wall-degrading enzymes, antibiosis, VOCs, and induced systemic resistance. Other research has also tested the biocontrol potential of this bacterial strain against various fungal and oomycete pathogens, with a successful degree of disease reduction reported ^37,61–63^.

## Conclusions

This study tested whether the bacterium *S. proteamaculans* and the mycoparasitic oomycete *Py. oligandrum* can counteract the devastating effects of *P. plurivora* on beech trees. Both *S. proteamaculans* and *Py. oligandrum* was found to reduce the prevalence of disease associated with *P. plurivora* during both *in vitro* and *in planta* greenhouse experiments, with the results suggesting that synergism between different mechanisms of action, including hydrolytic enzymes, VOCs, competition for nutrients, and induction of systemic resistance, underlies the reduced prevalence of the disease. Therefore, these antagonists may hold great potential for use as an alternative strategy to effectively control the adverse impacts of *P. plurivora* in beech, especially when combined with other disease control strategies.

## Author Contributions

All of the authors listed contributed to this research work and approved the manuscript for publication.

## Acknowledgments

RRV acknowledges support from FORMAS (2019–01316), Carl Tryggers Stiftelse för Vetenskaplig Forskning (CTS 20:464), the Novo Nordisk Foundation (0074727), and the SLU Centre for Biological Control. LGB is supported by the Swedish Research Council Vetenskapsrådet (2023-05529) and the Swedish Research Council Formas (2019-00881). SCW acknowledges funding from the Scottish Government’s Rural and Environment Science and Analytical Services (RESAS) Division.

